# Prior information enhances tactile representation in primary somatosensory cortex

**DOI:** 10.1101/2022.10.10.511201

**Authors:** Pegah Kassraian, Finn Rabe, Nadja Enz, Marloes Maathuis, Nicole Wenderoth

## Abstract

Perception and adaptive decision making rely on the integration of incoming sensory input with prior knowledge or expectations. While tactile stimuli play a significant role in shaping our perception and decision making, if and how prior information modulates the representation of tactile stimuli in early somatosensory cortices is largely unknown. Here, we employed functional magnetic resonance imaging (fMRI) and a vibrotactile detection paradigm to study the effect of prior information on tactile perception and tactile stimulus representation in early somatosensory areas. The supra-voxel somatotopic organization of early somatosensory areas allowed us to assess the effect of prior information on finger-specific representations. We found that vibrotactile stimuli congruent with expectations are associated with improved vibrotactile detection performance and a decrease of the mean blood-oxygen-level-dependent (BOLD) signal in the contralateral primary somatosensory cortex (S1). Concurrently, finger-specific activity associated with anticipated vibrotactile stimulation revealed higher multivariate decoding accuracies and better alignment with S1’s somatotopic organization. In addition, we observed that prior information induced somatotopically organized activity in contralateral S1 even before tactile stimulation onset. The accuracy of the multivariate decoding of stimulus-specific expectations was therefore strongly associated with upcoming behavioral detection performance. Thus, our results reveal a role for S1 in the integration of upcoming tactile stimuli with prior information based on its somatotopic organization as well as the presence of behaviorally relevant activity in S1 before stimulation onset.

## Introduction

A central mechanism facilitating perception and decision making is the integration of current sensory input with prior information or expectations. Previous research has shown that utilizing prior information can improve behavioral performance, leading to faster reaction times and improved stimulus detection (Kok et al., 2012; Richter and de Lange, 2019). The neural mechanisms underlying the integration of sensory input with prior information remain, however, largely unknown. Additionally, ongoing research is attempting to determine if and how prior information modulates the representation of stimuli in early sensory areas at the initial stages of sensory processing.

Most of the current knowledge on the neural mechanisms underlying the integration of prior information with sensory input stem from studies of the visual system. Functional magnetic resonance imaging (fMRI) in combination with visual detection paradigms has shown that anticipated visual stimuli are associated with a decrease in the blood oxygenation level dependent (BOLD) signal in early visual areas (Kok et al., 2012; Richter et al., 2018; Richter and de Lange, 2019). This result was initially surprising, as traditionally an increase of the BOLD signal had been linked to improved behavioral performance (Pessoa et al., 2003). The use of multivariate decoding analysis revealed, however, that representations of anticipated stimuli in the primary visual cortex can be decoded with higher accuracy. Together, these findings suggest that an enhanced signal-to-noise ratio during early sensory processing might underlie behavioral improvements when stimuli are anticipated (Kok et al., 2012, 2014, 2017; Richter and de Lange, 2019).

While touch is the first sense to develop prenatally (Reissland et al., 2014) and tactile perception is crucial in shaping decision making (Hernandez et al., 2000; Luna et al., 2005), it is largely unknown if the same mechanisms are at play in early somatosensory areas as relevant studies have remained scarce. Traditionally, early somatosensory regions have been thought to chiefly process afferent tactile stimuli. Recent work has demonstrated, however, that early somatosensory regions, such as the primary somatosensory cortex (S1), can be activated even in the absence of incoming sensory stimulation, for instance during touch observation, tactile imagery or during motor planning (Kuehn et al., 2014; de Borst et al., 2017, 2019; Gale et al., 2021; Kikkert et al., 2021; Ariani et al., 2022). Other work has shown that the history of vibrotactile stimulus presentation can bias the neural activity in brain regions related to encoding and memory (Preuschhof et al., 2010). Whether the expectation of tactile stimuli induces stimulus-specific activity patterns, if this activity is somatotopically organized, and if it is relevant for behavioral performance is, however, largely unknown. This is an important research question, as it has been hypothesized that such activity might influence the subsequent processing of anticipated stimuli (Kok et al., 2017; Blom et al., 2020; Aitken et al., 2020).

Here, we assessed if and how prior information modulates vibrotactile stimulus representations in early somatosensory regions both during and before vibrotactile stimulation. To this end, we employed fMRI and a vibrotactile detection paradigm where participants had to discriminate between weak, near-threshold vibrotactile stimulation of the ring finger versus the thumb after receiving probabilistic prior information on where to expect the stimulation. The vibrotactile stimulation intensities were set around the participants’ detection thresholds to reinforce the use of probabilistic prior information (Chambers et al., 2017; de Lange et al., 2018). The functional organization of early somatosensory areas in the form of somatotopic maps that stretch over a range of voxels allowed us to assess finger-specific modulation of vibrotactile stimulus representation by prior information.

We found that participants could detect vibrotactile stimulation congruent with their expectations more accurately and observed a decrease of the mean BOLD signal in contralateral S1 for these stimuli. Furthermore, ring finger versus thumb representations congruent with participants’ expectations were associated with higher linear decoding accuracies. Decoding accuracy and behavioral performance revealed a significant association, demonstrating the behavioral relevance of the precision by which congruent stimuli are encoded in contralateral S1. In addition, whole-brain searchlight decoding revealed that voxels assigned with higher decoding accuracies were more proximal to peak voxels determined by a functional localizer and univariate analysis.

Our paradigm allowed us to compare stimulus-specific activity patterns elicited by afferent stimulation to those elicited by the expectation of a vibrotactile stimulus. When investigating the period before vibrotactile stimulation onset, we found that a linear classifier could decode expectation of thumb versus ring finger stimulation based on the activity of contralateral S1. Crucially, we observed that BOLD activity during informative cue-stimulation intervals was increased within the region of interest (ROI) of the cued finger and concurrently suppressed within the ROI of the non-cued finger. Furthermore, the ability of a decoder to generalize between the activity patterns from afferent stimuli and activity patterns elicited by expectation demonstrated shared properties of the two representations. Finally, we observed that the decoding accuracy of ring finger and thumb cues were predictive of the upcoming behavioral performance, in line with the hypothesis that preparatory activity in early sensory areas might enhance perception and decision making.

## Results

### Prior information is associated with a decrease of the BOLD signal in contralateral S1

Twenty-five participants (15 males and 10 females) took part in a total of eight imaging runs over the course of two days (40 vibrotactile detection trials per run; Figure 1A; Materials and methods). Prior to each imaging run, the vibrotactile detection threshold of each participant’s ring finger and thumb were determined with an interleaved one-up one-down staircase (Figure 1B; Materials and methods).

**Figure 1.**
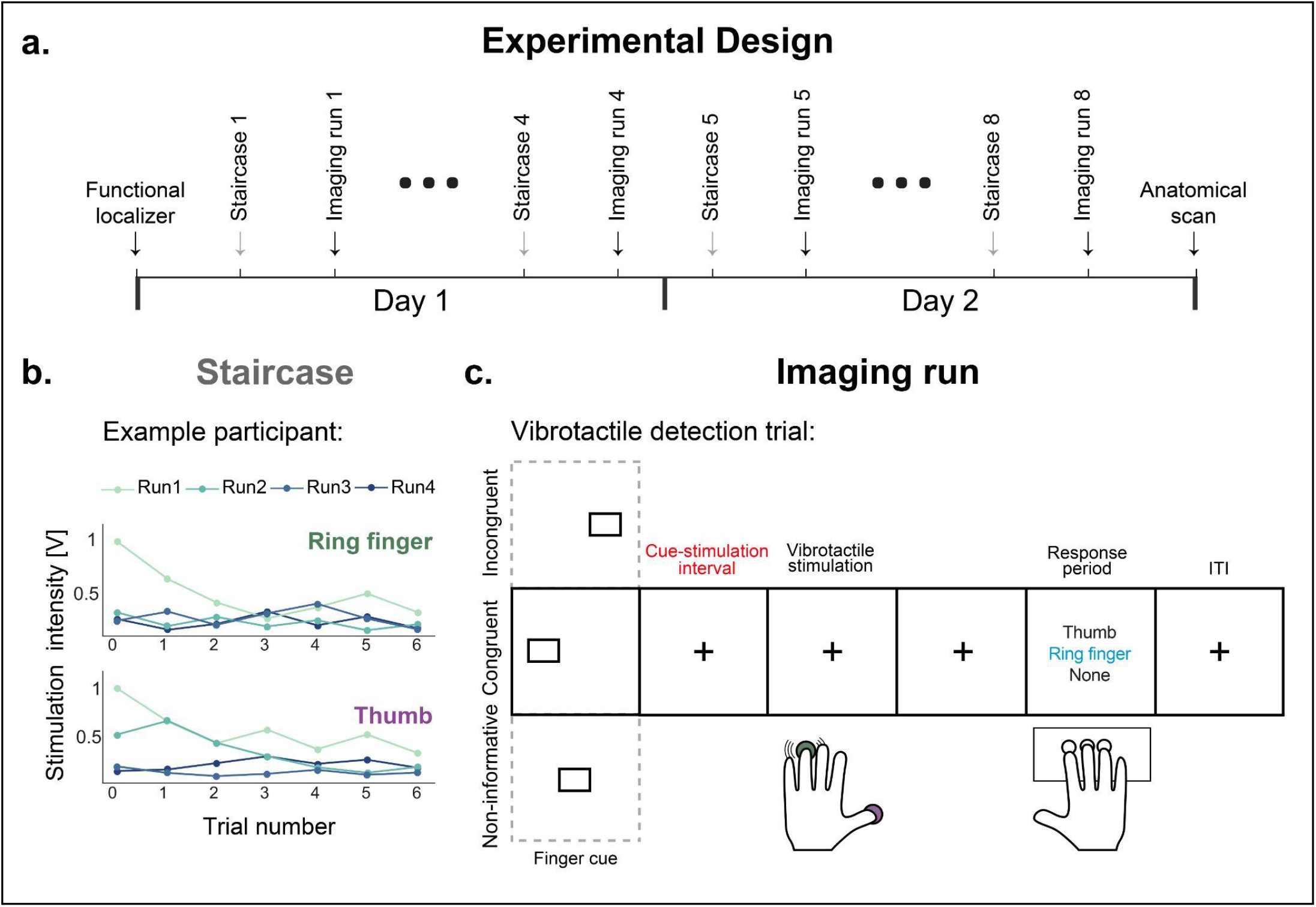
Experimental paradigm. **a.)** Participants took part in a total of 8 fMRI imaging runs on two consecutive days where they performed 40 vibrotactile detection tasks per run. Imaging runs were preceded by the acquisition of functional localizers on Day 1 and followed by the acquisition of structural MRI scans on Day 2. **b**.) Vibrotactile stimulation intensities of each imaging run were sampled around the vibrotactile detection threshold of the respective finger, determined by an interleaved staircase prior to the imaging run. Shown are the Day 1 results of an example participant. **c**.) Task structure of a vibrotactile detection trial: Each trial within an imaging run started with either a ring finger cue (square on the left), thumb cue (square on the right) or a non-informative cue (square in the middle of screen), followed by vibrotactile stimulation of either the ring finger (80% likely after ring finger cue; 50% likely after non-informative cue) or thumb (80% likely after thumb cue; 50% likely after non-informative cue) of the participant’s left hand. During the response period participants indicated where they had perceived the stimulation (‘Thumb’, ‘Ring finger,’ or ‘None’ if no stimulation was detected). Illustrated is an example congruent trial (ring finger cue followed by ring finger stimulation; dashed squares show possible alternatives). ‘+’: Fixation cross; ITI: Intertrial interval.

Each vibrotactile detection trial consisted of three parts (Figure 1C). First, a ‘finger cue’ which provided probabilistic prior information on upcoming stimulation location was shown, as described in the next paragraph. Second, there was vibrotactile stimulation of either the ring finger or the thumb of the participant’s left hand. Stimulation intensities were sampled around each participant’s detection threshold for the respective finger (Materials and methods). The use of weak vibrotactile stimulation with near detection threshold intensity was meant to reinforce the use of prior information. Third, there was a response period in which participants had to choose one of the three response options: ‘Thumb’, ‘Ring finger’, or ‘None’ if no stimulation was perceived (Figure 1C). The arrangement of the three response options on the screen was randomized between trials to preclude preparatory motor movements. The three parts of the trial were separated by intervals of variable lengths during which subjects were instructed to fixate on the displayed cross.

The finger cue in the form of a square was displayed on different parts of the screen: left, center, or right. Subjects were informed that a square on the left indicated that there was an 80% probability vibrotactile stimulation of the ring finger would follow (ring finger cue), a square on the right indicated that there was an 80% probability that stimulation of the thumb would follow (thumb cue), and a square in the middle indicated that there was an equally likely probability that stimulation of the ring finger or thumb would follow (non-informative cue). The ring finger and thumb were selected since somatosensory representations of these two digits can be dissociated particularly well with both univariate and multivariate methods (Martuzzi et al., 2012; Ejaz et al., 2015). The display location of the finger cue was spatially consistent with the position of the cued finger of participants’ left hand, thus ensuring an intuitive mapping. We refer to the experimental condition where the finger cue was followed by stimulation of the indicated finger as ‘congruent’ (i.e., ring finger cue and stimulation of ring finger or thumb cue and stimulation of thumb), as ‘incongruent’ when followed by stimulation of the other finger (i.e., ring finger cue and stimulation of thumb or thumb cue and stimulation of ring finger) and else as ‘non-informative’.

We observed that vibrotactile detection performance was enhanced for the congruent condition (70.2±4.81%) when compared to the non-informative (61.05±6.8%) and the incongruent experimental conditions (56.02±6.7%, mean±SEM; Figure 2A). These results were also reflected in the psychometric curves where a left shift was apparent for the congruent condition, indicating a decrease in vibrotactile detection threshold (Supplementary Figure 1). Stimulation intensities were matched between the experimental conditions (Materials and methods and Supplementary Table 1), thus ensuring that differences in vibrotactile detection performance and BOLD activity levels were due to differences in prior information available to participants.

**Figure 2.**
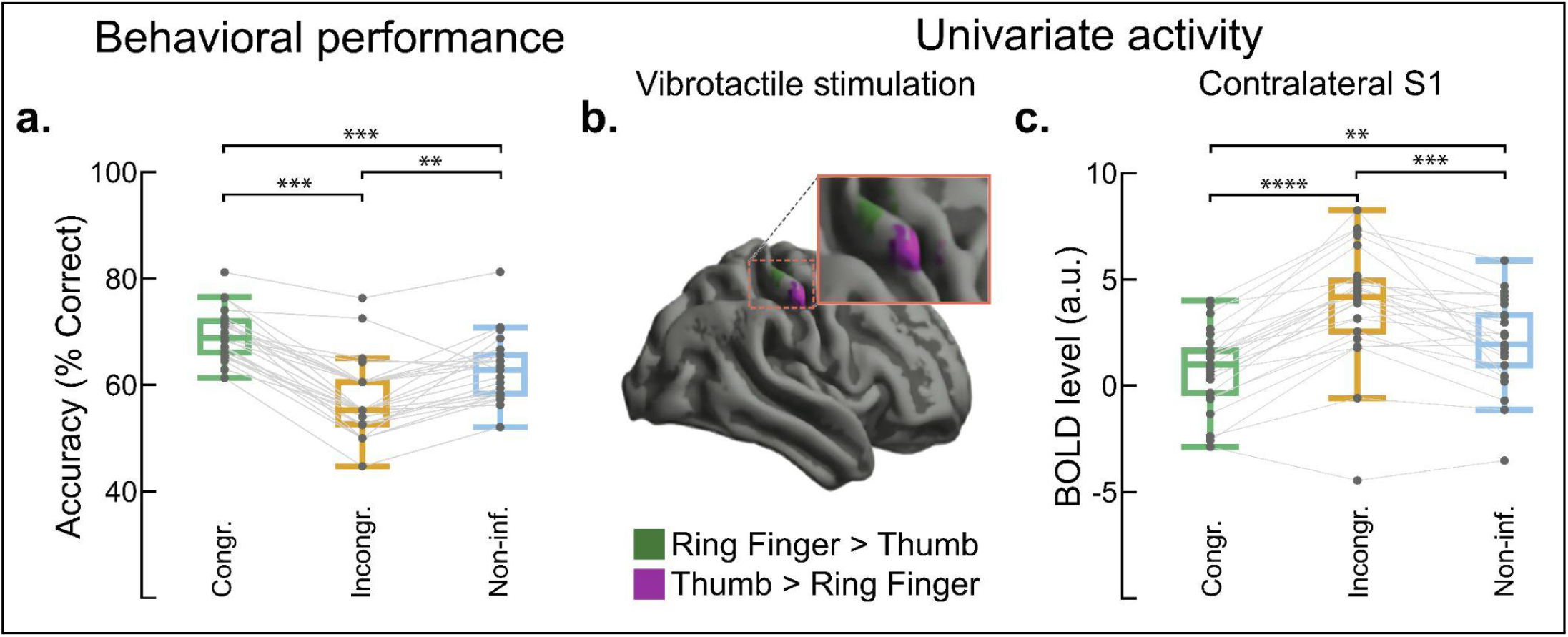
Effect of prior information on behavioral detection performance and BOLD signal. **a.)** Behavioral detection accuracies are enhanced for congruent (70.2±4.81%; mean±SEM) as compared to incongruent (56.02±6.7%) and non-informative trials (61.05±6.8%; one-way repeated measures ANOVA for Experimental condition: *F*_*2,48*_ = 87.839, p<0.001, η^2^ = 0.416). **b.)** Group-level GLM contrasts reveal distinct clusters of voxels in contralateral S1 for vibrotactile stimulation of ring finger versus thumb. Ring finger > Thumb: peak t-value = 5.182 (MNI coordinates: 42, -24, 60; cluster of 73 voxels); Thumb > Ring finger: peak t-value = 8.451 (MNI coordinates: 54, -14, 50; cluster of 128 voxels); p<0.05, familywise error (FWE) corrected. **c.)** The mean BOLD signal in contralateral S1 is dampened for congruent (0.796±0.368) as compared to incongruent (3.887±0.539) and non-informative trials (2.029±0.415; mean±SEM in a.u.: arbitrary units; one-way repeated measures ANOVA for Experimental condition: *F*_*2,48*_ = 44.466, p<0.001, η^2^ = 0.252). Pairwise t-tests with post-hoc Bonferroni correction: *** p<0.001, **** p<0.0001. Single-participant-results are visualized by gray lines (N=25 throughout all Figures).

To assess the BOLD signal associated with the vibrotactile stimulation of each finger, we performed a univariate contrast of ring finger versus thumb stimulation with a general linear model (GLM; Materials and methods). The contrast revealed two distinct clusters of voxels, one coinciding with the somatosensory ring finger ROI and one with the somatosensory thumb ROI, as determined by an independent localizer where supra-threshold stimulation was applied to participants ring finger or thumb (Figure 2B and Supplementary Table 2). The location of the clusters was in line with somatotopic finger representations in contralateral S1 as identified by previous fMRI studies (Martuzzi et al., 2012; Akselrod et al., 2021). Next, we compared the magnitude of the BOLD signal in S1 during vibrotactile stimulation events separately for the three different experimental conditions. Studies of the visual system have previously observed a decrease of the BOLD signal when sensory stimuli were anticipated, i.e. congruent with participants’ expectations as compared to the BOLD signal of ‘incongruent’ sensory stimuli (Kok et al., 2012; Richter and de Lange, 2019). Our results confirmed these findings: The average BOLD signal within contralateral S1 evoked by vibrotactile stimulation was significantly lower for the congruent as compared to the incongruent and non-informative experimental conditions (Figure 2C). No differences between the BOLD signal evoked by the three experimental conditions were observed in the ipsilateral S1 (Supplementary Figure 2).

### Prior information enhances stimulus representation in contralateral S1

Due to their supra-voxel somatotopic organization, early somatosensory regions are particularly well suited to study stimulus-specific modulation of activity through top-down signals. Here, we used linear decoding to classify the stimulated finger (ring finger versus thumb) separately for the three experimental conditions based on beta estimates obtained from a first-level GLM within anatomically defined ROIs (ROI-based decoding). We found that decoding accuracies for contralateral S1 were enhanced for congruent vibrotactile stimuli (63.067±1.311%, p<0.001) when compared to non-informative (56.51±1.419%, p<0.001) or incongruent experimental conditions (49.872±1.003%, p=0.4; all values mean±SEM; Figure 3A). These results were in line with previous studies demonstrating that anticipated visual stimuli are associated with higher multivariate decoding accuracies despite the observed concurrent decrease of the mean BOLD signal in early sensory areas (Kok et al., 2012; Richter and de Lange, 2019). Interestingly, decoding accuracies did not significantly differ from chance levels in ipsilateral and contralateral secondary somatosensory cortices. Decoding accuracies were moderately above chance for the contralateral primary motor cortex (Supplementary Figure 3A).

**Figure 3.**
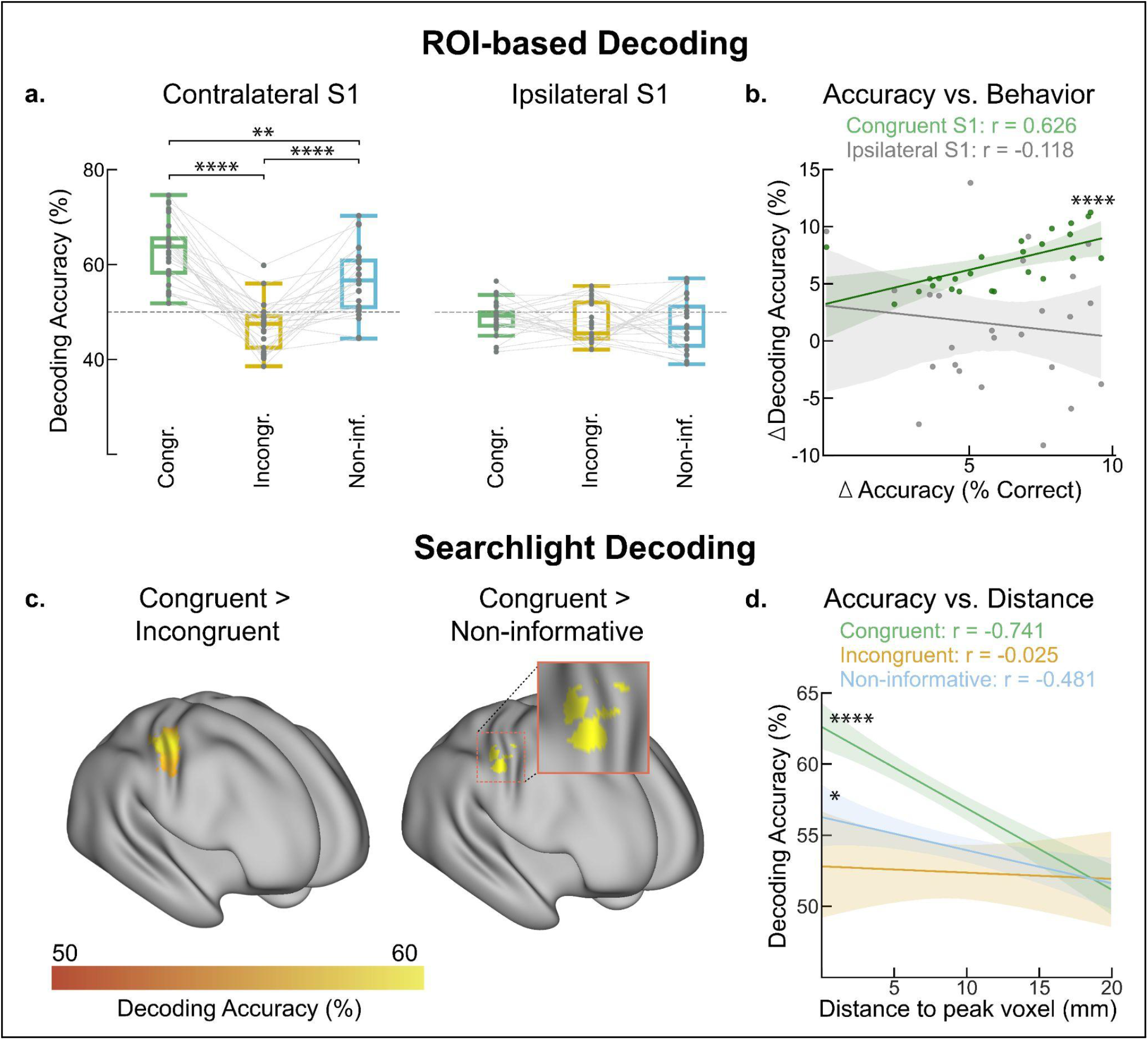
Multivariate decoding of vibrotactile ring finger versus thumb stimulation. **a.)** Contralateral S1 ROI-decoding accuracies are enhanced for congruent (63.067±1.311%, p<0.001) as compared to incongruent (49.872±1.003%, p=0.4) and non-informative trials (56.51±1.419%, p<0.001; mean±SEM; p-value calculation against empirical chance values from shuffled data). Decoding accuracies for ipsilateral S1 were not significantly greater than chance (p>0.4). Repeated-measures two-way ANOVA: Interaction Experimental condition x ROI, *F*_*2,96*_ *=* 17.847, p<0.0001, η^2^ = 0.294. Pairwise t-tests with post-hoc Bonferroni correction: **** p<0.0001. Single-participant-results are visualized by gray lines. **b.)** Improvements of behavioral detection accuracies (x-axis) and ROI-based decoding accuracies (y-axis) for congruent trials are strongly correlated for contralateral S1 (Spearman’s r, **** p<0.001, robust regression slope: 0.594; shaded area with 95% CI) but not for ipsilateral S1 (Spearman’s r, p=0.575, robust regression slope: -0.272 with shaded area: 95% CI). **c.)** Group-level searchlight maps reveal voxels throughout S1 and adjacent M1 with significantly higher decoding accuracies for the Congruent > Incongruent (left) comparison. The Congruent > Non-informative comparison (right), revealed two distinct clusters proximal to univariate ring finger and thumb ROIs (one-sided t-tests with p<0.01 and FWE-correction at the cluster-level with p<0.01). **d.)** Contralateral S1 searchlight decoding accuracies assigned to a voxel (y-axis) as a function of its Euclidean distance to the peak BOLD voxel (x-axis; MNI coordinates of peak BOLD voxel obtained by a functional localizer) reveal a strong correlation for congruent trials (Spearman’s r: **** p<0.001, * p<0.05, shaded area: 95% CI around robust regression line). Results averaged across ring finger and thumb, for a detailed discussion see Materials and methods.

To assess the behavioral relevance of the decoding results, we tested if the improvement in behavioral detection performance and decoding performance from the non-informative to the congruent experimental conditions are correlated (Figure 3B). This analysis revealed a strong correlation between behavioral detection performance and decoding performance in contralateral S1, demonstrating the behavioral relevance of the precision by which congruent stimuli are encoded in contralateral S1. In contrast, no such correlation was observed for the contralateral M1 region (Supplementary Figure 3B).

Next, we employed a searchlight decoding approach where a linear classifier with the same decoding scheme (ring finger versus thumb decoding; beta estimates from a first-level GLM) was applied one-by-one to each voxel and surrounding voxels (i.e., with a searchlight radius = 1; Kriegeskorte et al., 2006). This allowed us to obtain spatially more fine-grained accuracy maps across the whole brain. Group-level results of the searchlight decoding revealed clusters of voxels with significant decoding accuracies only for the congruent condition (Figure 3C). The largest cluster of voxels was located in the contralateral S1, with a subset of voxels localized in the contralateral primary motor cortex (787 voxels total with 62.6±4.3% decoding accuracy, mean±SEM). Significantly higher decoding accuracies between congruent versus incongruent or congruent versus non-informative experimental conditions were exclusively located in contralateral S1, proximal to the thumb and ring finger’s peak BOLD voxels (Figure 3C, Supplementary Table 3). Crucially, we observed in contralateral S1 a strong correlation between searchlight decoding accuracies of the congruent condition associated with a voxel and the distance of the voxel to the peak BOLD voxel of the respective finger (Figure 3D). This relationship was weaker for the non-informative experimental condition and not significant for the incongruent experimental condition.

Taken together, our results suggest that finger representations of congruent vibrotactile stimulations are associated with higher multivariate information content, are more aligned with the somatotopic organization of contralateral S1, and that the enhanced representation of these stimuli is strongly associated with behavioral detection performance.

### Distinct representations of ring finger and thumb cues in contralateral S1 prior to vibrotactile stimulation

Recent studies have shown that the primary somatosensory cortex can be activated even in the absence of incoming tactile stimulation, for instance during touch observation or tactile imagery (de Borst et al. 2017, 2019; Kuehn et al., 2019). It is, however, not known if expectation elicits somatotopic activity in S1 before stimulus onset, how such activity compares to activity evoked by afferent vibrotactile stimulation, and what the behavioral relevance of such activity would be. To address these questions, we analyzed the evoked BOLD activity during a variable interval following finger cue display and before vibrotactile stimulation onset (‘cue-stimulation interval’, Figure 1C). We categorized these periods as ‘informative’ if they followed a ring finger or a thumb cue and as ‘non-informative’ if they followed an non-informative finger cue (Figure 4A). The non-informative experimental condition served as a control and was divided into two groups based on the succeeding vibrotactile stimulation site (ring finger versus thumb stimulation).

**Figure 4.**
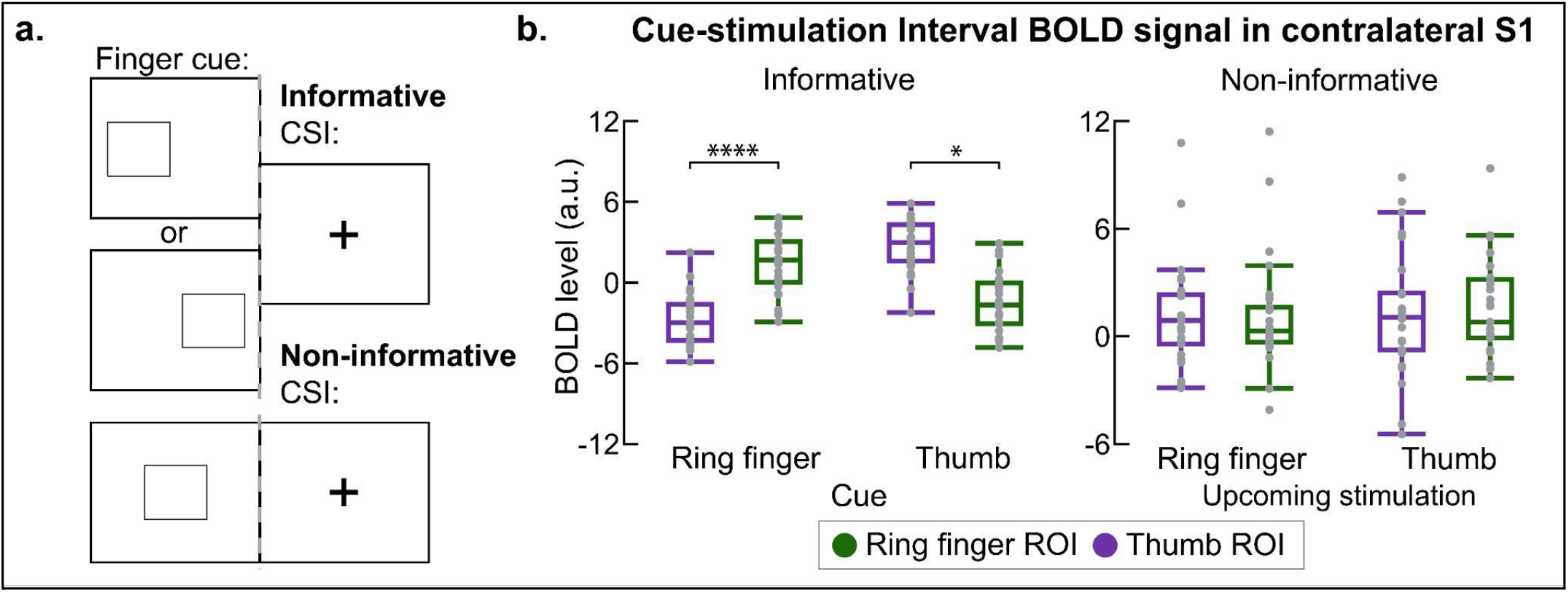
Somatotopic modulation of the BOLD signal during the Cue-stimulation interval (CSI). **a.)** The period between finger cue display and vibrotactile stimulation (CSI: ‘Cue-stimulation interval’; Figure 1C) was categorized as ‘informative’ when following a ring finger cue (square on left of screen) or a thumb cue (square on right of screen). Non-informative cue-stimulation intervals (square in the middle of screen) served as a control and were divided into two groups according to the following vibrotactile stimulation (ring finger vs. thumb). **b.)** After display of a finger cue, the BOLD signal was greater within the respective ROI (ring finger vs. thumb ROI) than within the ROI associated with the non-cued finger. No such modulation of BOLD activity could be observed after display of a non-informative finger cue. Ring finger and thumb ROIs were extracted for each participant from independent functional localizers (Materials and methods). A 2×2×2 way repeated measures ANOVA with CSI type x ROI x Cue condition revealed significant differences between the BOLD signal associated with ring finger vs. thumb cue for the informative CSI (p<0.01, t>4.98) but not for the non-informative CSI (p>0.258, t>0.0508). Pairwise t-tests with post-hoc Bonferroni correction: * p<0.01, ** *p<0.0001.

Notably, we observed that during informative cue-stimulation intervals the BOLD signal was increased within the ROI of the cued finger and concurrently suppressed within the ROI of the non-cued finger, thus revealing somatotopic organization (Figure 4B, left panel). In contrast, the BOLD signal during the non-informative cue-stimulation interval did not reveal any such organization (Figure 4B, right panel).

We next used a linear classifier to decode the cued finger (ring finger versus thumb cue) based on first-level GLM beta estimates obtained from the informative cue-stimulation interval, and, as a control, from beta estimates obtained from the non-informative cue-stimulation interval (Materials and methods). We found that decoding accuracies of the informative cue-stimulation interval were significantly higher than chance level in contralateral but not in ipsilateral S1 (Figure 5A). In contrast, it was not possible to decode the upcoming stimulation site from the non-informative cue-stimulation interval in any of the ROIs.

**Figure 5.**
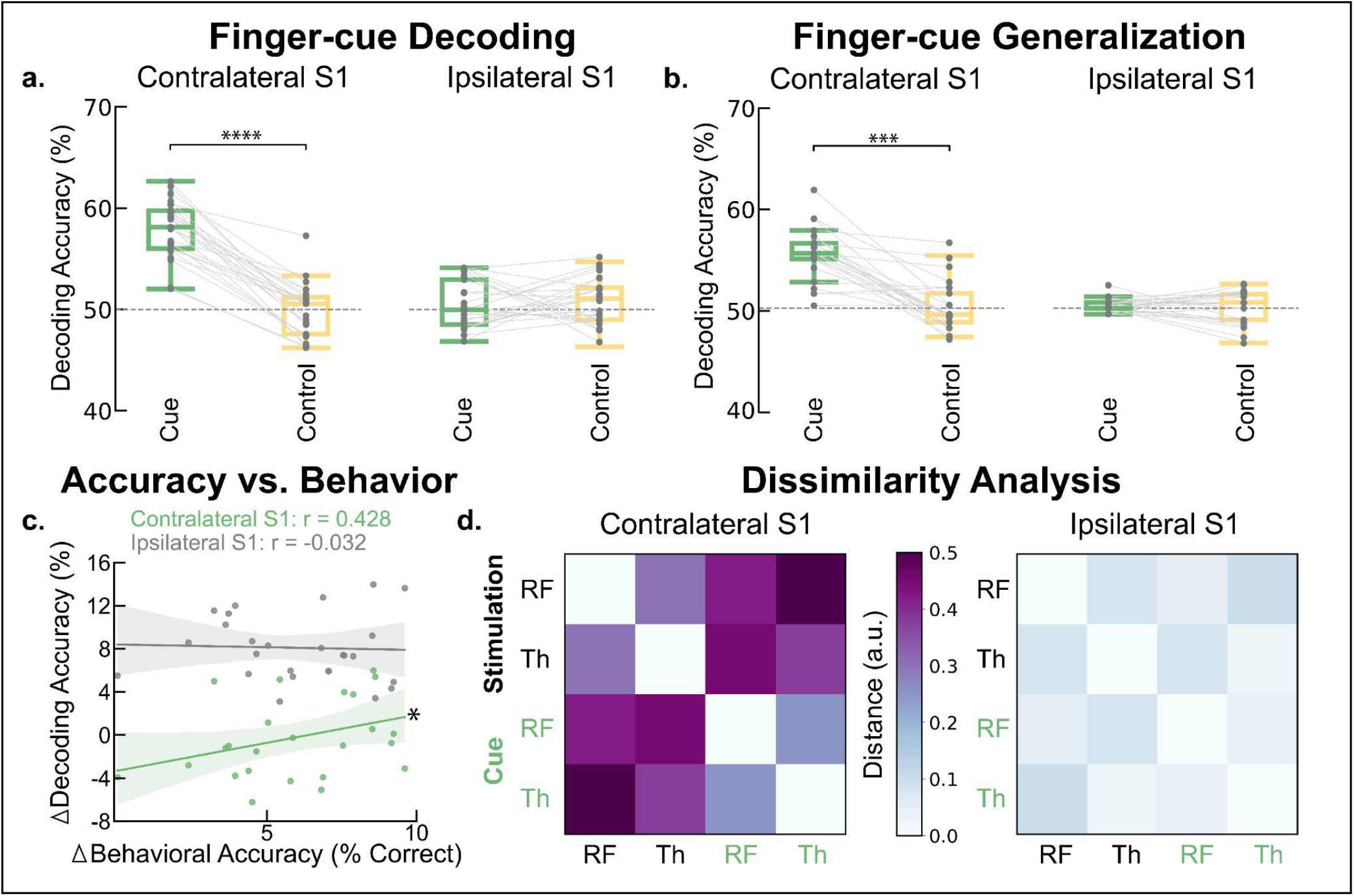
Decoding of ring finger versus thumb cue during the Cue-stimulation interval (CSI). **a.)** Accuracies of ring finger vs. thumb cue decoding are greater than chance for contralateral S1 during the informative but not during the non-informative CSI (57.327±2.8%, mean±SEM; p<0.0001 vs. 49.024±5.501, p=0.72). Decoding accuracies are not greater than chance for either condition for ipsilateral S1 (49.451±4.712 vs. 50.122±2.514%; p>0.62). Two-way repeated measures ANOVA: Interaction CSI type x ROI, F_1,24_ = 63.26, p<0.0001, η^2^ = 0.423. **b.)** Results from decoders trained on the CSI (ring finger vs. thumb cue decoding) and tested on vibrotactile ring finger vs. thumb stimulation trials and vice versa. Decoding accuracies were significantly greater than chance for the informative but not for the non-informative CSI for contralateral S1 (accuracies averaged over both train-test schemes: 56.921±3.754% vs. 50.191±1.207%, p<0.0001), but not for ipsilateral S1 (48.934±4.813%, p>0.7 vs. 49.325±3.204%, p>0.6). Two-way repeated measures ANOVA: Interaction CSI type x ROI, F_1,24_ = 35.385, p<0.0001, η^2^ = 0.308. Pairwise t-tests with post-hoc Bonferroni correction: **** p<0.0001. Single-participant-results are visualized by gray lines. **c.)** Improvement of behavioral detection accuracies (x-axis; from non-informative CSI to congruent stimulation trials) and ROI-based decoding accuracies (y-axis) for congruent trials are correlated for contralateral S1 (Spearman’s r, * p=0.039, robust regression slope: 0.539 with shaded area: 95% CI) but not for ipsilateral S1 (Spearman’s r, p=0.728, robust regression slope: -0.138 with shaded area: 95% CI). **d.)** Representational dissimilarity matrices visualizing the average cross-validated Mahalanobis distances (Walther et al., 2016) between ring finger (RF) and thumb (Th) stimulation (black labels) and ring finger and thumb cues during the CSI (green labels) for the contralateral (left) and the ipsilateral S1 (right).

We then asked if a linear decoder trained to discriminate between the anticipation of ring finger versus thumb stimulation would generalize to vibrotactile stimulation trials and vice versa. A decoder trained on informative cue-stimulation intervals and tested on vibrotactile stimulation periods and vice versa revealed significant accuracies for contralateral but not for ipsilateral S1 (Figure 5B). Training the decoder on non-informative cue-stimulation intervals resulted in chance level-accuracies for both ROIs, thus supporting the hypothesis that the anticipation of vibrotactile stimulation was required to drive informative activity during this period.

We then compared the improvement of decoding performance from non-informative to informative cue-stimulation intervals to the behavioral improvement from non-informative to congruent vibrotactile stimulation trials (Figure 5C). This comparison revealed a significant correlation in contralateral but not in ipsilateral S1, demonstrating the behavioral relevance of S1 activity even prior to vibrotactile stimulus onset, although the association was weaker than the association found during vibrotactile stimulation (Figure 3B). Consistent with these results, we found that Mahalanobis distances were greatest between the activity evoked by vibrotactile stimulation of ring finger vs. thumb. In addition, activity evoked by a finger cue bore greater similarity to activity evoked by the vibrotactile stimulation of the corresponding finger (Figure 5D).

In summary, we observed that expectation of vibrotactile stimulation to a finger elicits somatotopic activity in contralateral S1, that this activity allows the decoding of finger-specific anticipation, and that it bears similarity to the activity elicited by afferent vibrotactile stimulation in contralateral S1.

## Discussion

The integration of afferent sensory information with prior expectations facilitates perception and decision making, for instance when sensory input is weak or ambiguous. In this study, participants received weak vibrotactile stimulation to either the ring finger or thumb preceded by a probabilistic finger cue, and had to indicate on which finger they had perceived the stimulation. We found that vibrotactile stimulation congruent with the participants’ expectations was associated with enhanced detection performance and observed a concurrent decrease of the BOLD signal in contralateral S1. This result is consistent with studies of the visual system demonstrating that anticipated visual stimuli are associated with enhanced detection performance and a concurrent dampening of the BOLD signal in primary visual cortex.

Multivariate decoding of the stimulated finger (ring finger versus thumb) revealed improved decoding accuracies for vibrotactile stimuli congruent with participants’ expectations when compared to stimuli which were incongruent or followed an non-informative finger cue. Importantly, we observed a significant association between improvements of detection performance and decoding accuracy for anticipated stimuli, demonstrating the behavioral relevance of the precision by which anticipated vibrotactile stimuli were encoded in S1. In contrast, we did not find linear decoding accuracies to be significantly higher than chance level in the ipsilateral and contralateral S2 regions. This is consistent with findings of neuroimaging studies demonstrating that individual finger representations cannot be resolved in S2 with 3T or even with 7T fMRI (Ruben et al., 2001; Sanchez-Panchuelo et al., 2018) and with primate electrophysiology showing that S2 neurons have larger and often bilateral receptive fields with a less fine-grained somatotopic organization (Fitzgerald et al., 2006). In light of findings that top-down signals such as attention can modulate activity both in primary and secondary somatosensory cortices (Johansen-Berg et al., 2000), this might indicate that a modulation of S2 activity by top-down signals relies less strongly on a somatotopic organization.

In addition, we found decoding accuracies in the ipsilateral and contralateral primary motor cortex to be only moderately above chance and not associated with behavioral detection performance. Whole brain searchlight decoding confirmed that the effects of prior information were largely confined to S1, with peak decoding accuracies being located proximal to univariate peak voxels as determined by an independent functional localizer. Thus, our results suggest that the behavioral improvements resulting from prior information are associated with higher separability of somatotopic stimulus representations in contralateral S1. These results provide evidence for the integration of prior information during early somatosensory processing stages, where the activity of a select neural population could be ‘tuned’ for the anticipated tactile stimulus.

The finding that prior information can modulate activity in early sensory areas that had been traditionally associated with the processing of afferent stimuli has prompted the question of whether top-down signals alone can activate these regions. Studies of the visual and auditory systems have revealed that expectation of a stimulus indeed increases baseline activity in the relevant early sensory areas, leading to the hypothesis that this allows for efficient perceptual processing (Bell et al., 2016; Kok et al., 2014, 2017; Blom et al., 2020). Two recent studies have provided converging evidence for the involvement of S1 in motor planning (Gale et al., 2021; Ariani et al., 2022), with one study demonstrating evidence for finger-specific activity in S1 during motor planning (Ariani et al., 2022). How stimulus-specific expectation modulates activity in early somatosensory areas, and, importantly, how such activity compares to somatosensory activity elicited during perception of afferent tactile stimuli has so far, however, not been studied.

When assessing the period before vibrotactile stimulation, we found that informative but not non-informative probabilistic finger cues induce anticipatory activity patterns in contralateral S1. Importantly, we observed finger-specific modulation of the BOLD signal during this period.

In addition, ring finger and thumb cues could be decoded from BOLD activity after an informative cue (ring finger versus thumb cue) and the performance of the linear decoder was related to the upcoming detection performance of the participants. In a subsequent analysis, we assessed the similarity between the activity patterns elicited by anticipation of a vibrotactile stimulus and the activity patterns associated with vibrotactile stimulation. Crucially, we found that a linear decoder trained during the anticipation of a vibrotactile stimulus could decode activity patterns from the associated vibrotactile stimulation with above-chance accuracy and vice versa. These findings are in line with the hypothesis that top-down modulation of activity in primary sensory cortices rely on similar neural mechanisms as those relevant for perception.

In summary, our results demonstrate that prior information can drive somatotopic activity in early somatosensory areas even in the absence of afferent tactile stimulation, and that this activity is task relevant. This finding adds to the recent literature demonstrating that S1 is involved in higher order processing of tactile information based on top-down signals (de Borst et al., 2017, 2019; Ariani et al., 2022), and that it does so based on its somatotopic organization (Ariani et al., 2022). Taken together, our results are in line with the processing of prior information during early stages of sensory processing, as suggested by inferential models of perception.

Activity in both early and late sensory areas has been associated with a range of top-down cognitive processes, such as mental imagery or feature-based attention (Liu et al., 2016; Koenig-Robert and Pearson, 2019; Blom et al., 2020). With respect to the somatosensory domain, recent studies have demonstrated for instance that both tactile imagery as well as motor planning evoke informative activity patterns in somatosensory areas (de Borst et al., 2017, 2019; Gale et al., 2021; Ariani et al., 2022). It would be of interest to further refine the quantification of the involved top-down processes. It is, for instance, not yet known to what extent motor planning involves mental imagery, and while it has been shown that motor preparation elicits somatotopic activity in S1 similar to the activity induced by motor execution, neural representations associated with mental imagery have only been found to allow for the dissociation of the imagery modality (tactile versus auditory) but not for stimulus-specific imagery content (de Borst et al., 2017). This note also extends to our study, where we cannot dissociate between the expectation of an upcoming vibrotactile stimulus and the respective mental imagery. Thus, the assessment of the differences between different categories of anticipatory neural activity both in somatosensory areas and areas associated with other stimulus modalities constitutes a promising future line of research.

## Author contributions

Conceptualization: PK, NW; Data collection: PK, NE; Methodology: PK, FR, MM, NW; Software and data analysis: PK, FR; Writing - original manuscript: PK, Writing - review and editing: PK, FR, MM, NW; Funding acquisition: PK, NW; Supervision: MM, NW.

## Acknowledgements

We thank the participants of the study, Daniel Woolley for feedback on the manuscript and initial support with the vibrotactile stimulation setup, members of the Wenderoth lab for participating in study pilots and Roger Luechinger for the technical support during fMRI data collection.

## Data availability

The data used to create the figures in this study can be found on Github https://github.com/Pegahka/PriorInformationS1

## Materials and methods

### Participants, Power, Ethics

25 healthy volunteers participated in the study (15 males, 10 females, age 26±4 years; mean±SD; all participants right-handed as assessed by the Edinburgh Handedness Inventory). The sample size is in line with sample sizes employed in the field (Kok et al., 2012, Ariani et al., 2022). Participants’ informed consent was obtained according to the Declaration of Helsinki prior to study onset. Ethical approval was granted by the Kantonale Ethikkommission Zürich (KEK-2015-0537).

### Experimental design

After familiarization in a mock-scanner, participants performed vibrotactile detection tasks in a functional magnetic resonance imagining (fMRI) scanner during 2 sessions on two subsequent days. Each fMRI imaging session consisted of four imaging runs. Each imaging run (∼10 minutes) consisted of 40 vibrotactile detection trials and was preceded by a psychophysical staircase procedure (∼7 minutes). In addition, day 1 began with functional localizer runs (2 runs of each 10 minutes) and a structural scan was obtained for each participant at the end of day 2 (∼7 minutes), resulting in a duration of around 1.5-2 hours for each fMRI imaging session.

### Vibrotactile stimulation

Custom-made MR compatible piezo-electric devices (Dancer Design, UK) were used to deliver stimulation to the fingertips of participants’ ring finger and thumb of the left hand. Stimulators were fixated on the fingers using Velcro straps to ensure good contact with the skin throughout the experiment without compromising blood supply. A pulsed sinusoid with 5 bursts per stimulation at a frequency of 50 Hz with 40 ms gaps and a duration of 3.8 seconds was employed for vibrotactile stimulation throughout the entire study. Visual input was displayed on a screen visible to participants through a mirror placed on the head coil. Stimulation and display were controlled by in-house scripts implemented in MATLAB (2018a, https://www.mathworks.com) using the Psychophysics Toolbox (Brainard, 1997, Pelli, 1997, Kleiner et al., 2007). Two interleaved one-up one-down staircases were employed prior to each imaging run, to determine vibrotactile detection thresholds separately for the ring finger and thumb of the left hand of each participant. Each staircase started at 1 Volt stimulation intensity and consisted of 7 trials per finger in a randomized order where participants had to indicate on which finger they perceived the stimulation. To ensure an experimental environment akin to the main experiment, staircases were conducted inside the scanner while echo-planar images (EPIs) were acquired, but the corresponding MR scans were later discarded. For the following imaging session vibrotactile stimuli were presented around the participants’ detection threshold with stimulation intensities sampled for each finger individually from a skew-normal distribution with a skewness of 0.5 and a kurtosis of 4, while the mode of the distribution was set to the vibrotactile detection threshold of that finger.

### Experimental imaging runs and behavioral tasks

Each vibrotactile detection trial began with the display of a finger cue (1s duration). Next, in the vibrotactile stimulation period (3.8s), participants received stimulation to either the thumb or ring finger of the left hand. Finally, there was a response period (2s) where participants indicated the perceived vibrotactile stimulation site: ‘Ring finger’, ‘Thumb’ or ‘None’, the latter included in case they could not detect any vibrotactile stimulation. Finger cue display, vibrotactile stimulation and Response period were separated by intervals of variable lengths sampled uniformly from the respective range (3.5-4.5s, 2.5-3.5s or 1-2s, respectively). Participants were instructed to focus on a fixation cross when displayed.

The finger cue (a square on a black background) appeared either on the left, right or center of the screen. A square on the left indicated that stimulation was likely (80% chance) to be applied to the left ring finger (ring finger cue), while a square on the right indicated that stimulation was likely (80% chance) to be applied to the thumb (thumb cue). A square in the middle indicated that stimulation to the thumb or ring finger was equally likely (non-informative cue). Note that the finger cue location was spatially consistent with the position of the cued finger of participant’s left hand, ensuring an intuitive mapping for the participants.

Trials where the stimulated finger matched the cue were classified as the ‘congruent’ experimental condition (thumb cue – stimulation of thumb; ring finger cue – stimulation of ring finger), while trials where the stimulated finger was incongruent to the cue were classified as belonging to the ‘incongruent’ experimental condition (thumb cue – stimulation of ring finger; ring finger cue – stimulation of thumb). The trials with the non-informative cue formed the ‘non-informative’ experimental condition. Of the total 320 trials, 172 fell into the congruent category, 44 into the incongruent category and 104 into the non-informative category. The same randomized order of experimental conditions (congruent, incongruent, non-informative) was employed for all participants, and it was ensured that each experimental condition contained the same number of ring finger and thumb trials. During the response period, participants could select from the three response options (“Ring finger”, “Thumb”, or “None”) with their right hand through key presses on a button box. The response options changed color upon selection, and their arrangement was randomized to prevent preparatory motor artifacts prior to the response period.

### Statistical analysis of the behavioral task

Behavioral detection accuracy (Figure 2A) was calculated as the fraction of correctly recognized stimulation sites over all vibrotactile stimulation events over the 8 runs. Statistical analyses were performed using the R statistical software (4.2.1; www.R-project.org). Post hoc Bonferroni corrected p-values for two-sided pairwise t-tests were also included.

We performed a robust regression analysis (with robustbase in R) to assess if the improvement of behavioral detection accuracy and the improvement of decoding accuracy observed for congruent stimuli were related (Figure 3B). To this end, the independent variable was calculated as the difference between behavioral accuracy for the congruent stimuli and the behavioral accuracy for non-informative stimuli for each subject. The dependent variable was calculated as the difference between decoding accuracy for the congruent stimuli and the decoding accuracy for non-informative stimuli for each subject either based on the BOLD signal in the contralateral S1 region (Figure 3C, left panel) or the ipsilateral S1 region (Figure 3C, right panel). Next we assessed the relationship between the improvement of behavioral detection accuracy for congruent stimuli and the decoding accuracy of the informative cue-stimulation interval to assess the behavioral relevance of the BOLD signal induced by a finger cue prior to vibrotactile stimulation. Here, the independent variable was calculated as the difference between the decoding accuracy for the informative cue-stimulation interval minus the decoding accuracy for the non-informative cue-stimulation interval for each subject. The robust regression analysis was again performed for both the contralateral S1 region and the ipsilateral S1 region (Figure 5C).

### Functional Localizer and finger ROI

A functional localizer was performed over 2 runs on day 1 prior to the start of the imaging runs of the main experiment. Fingers were stimulated with a clearly perceivable, suprathreshold vibrotactile stimuli (4 Volts stimulation intensity, 4s duration). Each finger was stimulated 15 times with variable interstimulus intervals between the stimulation periods. The marsbar ROI toolbox (http://marsbar.sourceforge.net) was used to extract ring finger and thumb region of interest masks for each participant based on the significant cluster of activity (p<0.05) in contralateral S1 from a vibrotactile stimulation > resting baseline contrast.

## fMRI data acquisition

Functional images were acquired on a Philips Ingenia 3T with a fifteen element head coil. The T2*-weighted gradient-echo transversal EPI sequences were acquired with a spatial resolution of 2.75×2.75×3.3 mm^3^, TR/TE = 2500ms/35ms and an epi-factor of 39 and a sense acceleration factor of 2. The 40 acquired slices with no interslice gap were oriented parallel to the ACPC line. Anatomical images were acquired with a 1 mm×1 mm×1 mm spatial resolution, TR/TE = 8.38 ms/3.94 ms and a sense acceleration factor of 1.5 and sagittal orientation.

### fMRI preprocessing and univariate analysis

The fMRI data was pre-processed using a standard pipeline (i.e., realignment, normalization, and smoothing) in FSL (v5.00; https://fsl.fmrib.ox.ac.uk/fsl/fslwiki/FSL). Realignment of the functional images was followed by skull stripping applied to the structural image with the brain extraction tool BET (Smith et al., 2002; Jenkinson et al., 2005). Structural images were segmented with FAST (Zhang et al., 2001) and the boundary based co-registration tool FLIRT (Jenkinson and Smith, 2001; Jenkinson et al., 2002) was then used to co-register functional images to the structural image. Motion correction was applied using MCFLIRT (Jenkinson et al., 2002). Spatial smoothing with a 3 mm full-width half maximum Gaussian kernel and a high-pass filter with a cut-off of 100 seconds were applied to the co-registered functional images. Finally, functional images were normalized to the MNI template space with FNIRT (Andersson et al., 2007). The preprocessed images were analyzed with a general linear model (GLM). We defined separate regressors for each vibrotactile stimulation and each cue-stimulation interval (period between finger cue display and vibrotactile stimulation). Finger cue display and the response period were each included as regressors of no interest, as were head motion parameters. Single-subject (first-level) beta estimates (regression coefficients) were obtained based on the double-gamma hemodynamic response function (canonical HRF) and its temporal derivatives. The first-level analysis resulted in activation images (beta maps) for each of the vibrotactile stimulations and each of the cue-stimulation intervals for each of the participants. Contrasts for main effects and interactions were defined at the first level and employed for the group-level (second-level) analysis. Results are reported at p < 0.05 and familywise error (FWE) corrected at the voxel level based on the Threshold-Free Cluster Enhancement (TFCE; Smith and Nichols, 2009). BOLD level analysis (Figure 2C) was performed by averaging the beta estimates for all stimulation events of an experimental condition over all voxels.

### Regions of interest definition

Regions of interest were extracted in MNI space with probabilistic cytoarchitectonic masks from the SPM Anatomy Toolbox (Version 3.0, Forschungszentrum Jülich GmbH, Eickhoff et al., 2005) and used for the ROI-based decoding analysis and BOLD level assessments. In particular, we extracted masks for the primary somatosensory cortices (S1; 3148 voxels), the secondary somatosensory cortices (S2; 2872 voxels), the primary motor cortex (M1; 4321 voxels); the primary visual cortex (V1; 1802 voxels) and for white matter regions (9321 voxels). We excluded voxels with a larger than 25% overlap on opposite sides of the central sulcus for a distinct analysis of S1 versus M1 (Ariani et al., 2022).

### Region of interest decoding

Samples for binary decoding were extracted within the relevant ROI from beta-maps obtained by the first-level GLM of each participant (vibrotactile finger stimulation > baseline or cue-stimulation intervals > baseline, resulting in 320 beta-maps used as decoding samples). The decoding analysis was implemented with a linear Support Vector Machine (SVM) with MATLAB’s fitcsvm function (https://www.mathworks.com/help/stats/fitcsvm.html). The linear SVM was employed for binary prediction of stimulated finger (ring finger or thumb; Figure 3A) or finger cue (Figure 5A). To match the number of voxels between different ROIs, 300 voxels were randomly subsampled from each ROI for each iteration of a 10-fold cross-validation. Reported decoding accuracies are cross-validation accuracies averaged over 1000 rounds. The SVM cost parameter was used in its default setting (i.e., *C*=1). The decoding was balanced by subsampling to obtain the same number of stimulation events per condition (e.g., same number of ring finger versus thumb samples for each of the three experimental conditions). Decoding labels were permuted to assess the significance of decoding accuracies and the empirical p-value was calculated as the fraction of decoding accuracies from the permuted dataset larger than accuracies obtained from the original dataset based on 1000 such permutations (Kassraian-Fard et al., 2016; Ojala and Garriga, 2009).

To assess the generalization performance of linear decoders (Figure 5B) across cue-stimulation intervals and vibrotactile stimulation trails, SVM decoders were trained using the parameters described and by using data from the cue-stimulation interval (ring finger vs. thumb cue decoding) and tested on vibrotactile stimulation trials (ring finger versus thumb stimulation decoding) and vice versa. The same procedure was implemented for non-informative cue-stimulation trials which were divided based on the upcoming stimulation site (ring finger versus thumb stimulation). Reported decoding accuracies are averages across the two train-test schemes.

### Whole-brain searchlight decoding

The searchlight analysis (Figure 3C and 3D) was implemented with the pyMVPA toolbox (Hanke et al., 2009) with a linear SVM (C=1) applied one by one to each voxel and its surrounding sphere with a radius of 1 voxel. The linear SVM was employed to predict stimulated fingers (ring finger or thumb) based on the 320 beta maps corresponding to the 320 stimulation events. Random subsampling of trials was implemented to ensure an equal representation of each of the two experimental conditions in train and test sets. Searchlight decoding resulted in a whole-brain accuracy map for each participant where the decoding accuracy was ascribed to the voxel in the center of the sphere. To obtain group-level results a one-tailed *t*-test across participants was performed against resting baseline or between experimental conditions. Group-level accuracy maps were thresholded and FWE corrected (p < 0.01). Searchlight accuracy maps were plotted with the connectome workbench (Figure 3C) (https://www.humanconnectome.org/software/connectome-workbench; Marcus et al., 2011).

Whole-brain searchlight decoding accuracy of a voxel (dependent variable) was compared to its Euclidean distance to the peak BOLD voxel associated with vibrotactile stimulation of ring finger or thumb (the independent variable; Figure 3D). This analysis was performed as follows: First, a searchlight accuracy map was obtained through binary classification of vibrotactile stimulation of each finger versus resting baseline, resulting in 50 searchlight accuracy maps (ring finger and thumb searchlight accuracy maps for 25 subjects). Then, we calculated for each voxel within the S1 region of a searchlight map its Euclidean distance to the MNI peak BOLD voxel associated with vibrotactile stimulation of the respective finger as obtained by an independent functional localizer. Finally, accuracies were averaged across both fingers for the respective distance bin. We performed a robust regression analysis (with robustbase in R) to determine the relationship between searchlight decoding accuracy and distance to peak BOLD voxel.

### Multivariate dissimilarity analysis

For the dissimilarity analysis between evoked activity from vibrotactile stimulation of ring finger and thumb or by ring finger and thumb cues (Figure 5D), we calculated the cross-validated Mahalanobis distance based on beta estimates from the first-level GLM of participants. The distance calculation was performed with a leave-one-run-out cross-validation schemed. Visualized results are averages over all voxels within the indicated ROI. Dissimilarity calculations were implemented by using the pyMVPA toolbox.

## Supplementary Material

**Supplementary Figure 1:**
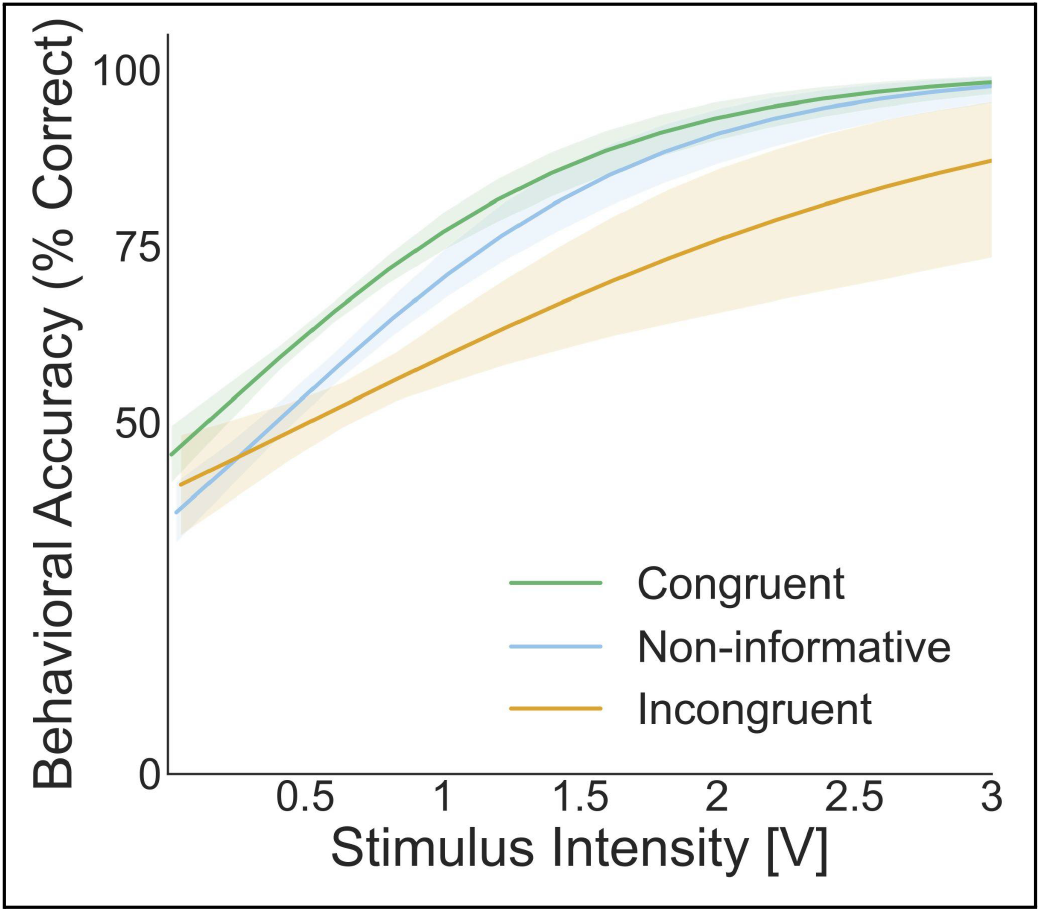
Psychometric curves for the vibrotactile detection task. Psychometric curves for vibrotactile stimulus detection (% correct) during a total of 320 trials over 8 imaging runs as a function of vibrotactile stimulation intensity (Volt). The psychophysical curve of the congruent condition is shifted to the left as compared to the psychophysical curves of the incongruent and the non-informative experimental conditions, indicative of a lower vibrotactile detection threshold for the former. Curves represent logistic fits to mean values across all participants (N=25), error bands around psychophysical curves represent the 95% bootstrap confidence intervals.

**Supplementary Table 1:** Average vibrotactile stimulation intensities for experimental condition and finger.

| Experimental Condition | Intensities Thumb [V] | Intensities Ring finger [V] |
| --- | --- | --- |
| Congruent | 0.94 ± 1.64 | 0.79 ± 1.5 |
| Incongruent | 1 ± 1.73 | 0.8 ± 1.67 |
| Non-informative | 0.98 ± 1.7 | 0.74 ± 1.35 |

Average vibrotactile stimulation intensities (mean± SEM) for 320 trials over 8 imaging runs for thumb and ring finger trials. A repeated measures two-way ANOVA with factors Finger x Experimental condition reveals no significant main effect of experimental condition for vibrotactile stimulation intensities ( $F_{1,2} = 0.55$ , $p = 0.58$ , $\eta^2 = 0.018$ ).

**Supplementary Figure 2:**
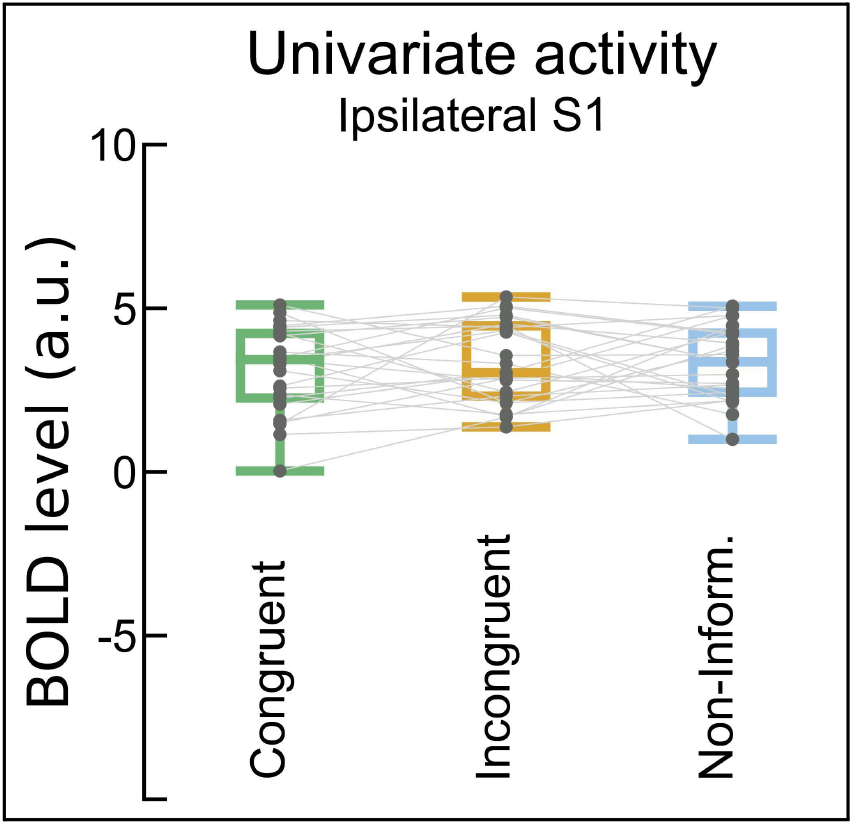
The ipsilateral S1 BOLD level is not modulated by prior information. The mean BOLD signal in ipsilateral S1 is not significantly different for congruent (3.229±0.249) vs. incongruent (3.355±0.33) and non-informative trials (3.743±0.261; a.u.: arbitrary units; one-way repeated measures ANOVA for Experimental condition: *F*_*2,48*_ = 1.789, p=0.71). Single-participant-results are visualized by gray lines (N=25).

**Supplementary Figure 3:**
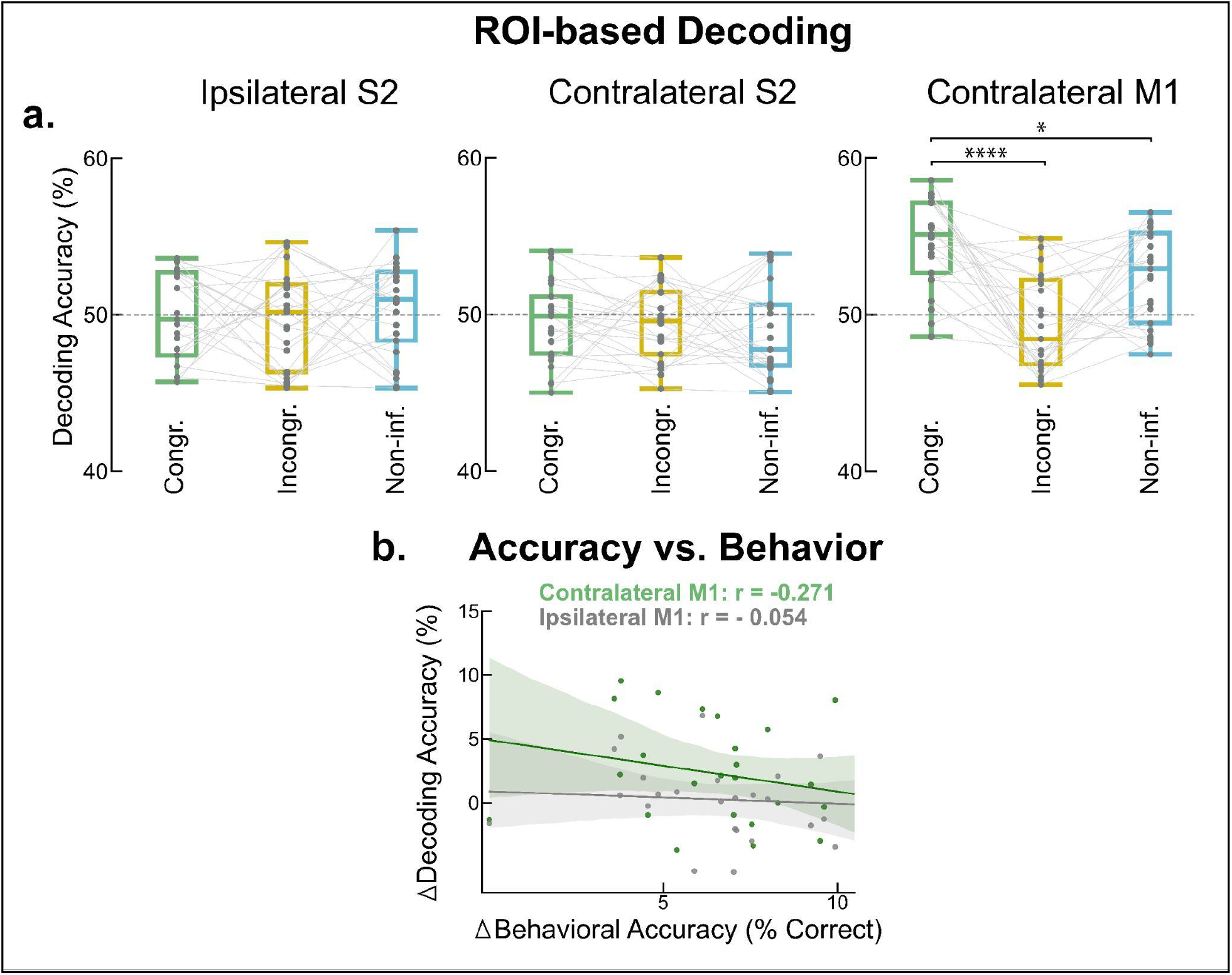
ROI-based decoding of vibrotactile stimulation of ring finger vs. thumb for secondary somatosensory and primary motor cortex. **a.** Left and center: Ipsilateral S2 ROI-decoding accuracies are not significantly different from chance level for any of the experimental conditions (Ipsilateral S2; congruent: 49.928±5.703%, incongruent: 49.726±6.392%, non-informative: 50.372±5.991% Contralateral S2; congruent: 49.527±5.382%, incongruent: 49.448±4.976%, non-informative: 48.89±5.832%, mean±SEM, p>0.5). Right: Decoding accuracies for contralateral M1 were significantly greater than chance for congruent (54.605±5.78%) and non-informative trials (52.368±6.11%), but not incongruent trials (49.368±4.928%). Repeated-measures two-way ANOVA: Interaction Experimental condition x ROI, *F*_*2,96*_ = 9.452, p<0.0001, η^2^ = 0.311. Pairwise t-tests with post-hoc Bonferroni correction: * p<0.05; **** p<0.0001. Single-participant-results are visualized by gray lines. **b.** Improvements of behavioral detection accuracies (x-axis) and ROI-based decoding accuracies (y-axis) are not correlated for contralateral (Spearman’s r, p=0.181, robust regression slope: 0.594 with shaded area: 95% CI) or ipsilateral M1 (Spearman’s r, p=0.71, robust regression slope: -0.272 with shaded area: 95% CI).

**Supplementary Table 2:** Peak t-values associated with vibrotactile stimulation of ring finger versus thumb.

|  | *Condition* | *BA* | *Hemisphere* | *x* | *y* | *z* | *Peak t-values* |
| --- | --- | --- | --- | --- | --- | --- | --- |
| **Peak t-values** | Th. > Ringf. | Postc. Gyr. (SI) 3b | R | 54 | -14 | 50 | 8.451 |
|  | Ringf. > Th. | Postc. Gyr. (SI) 1 | R | 42 | -24 | 60 | 5.182 |

FWE correction with random field theory, $p < 0.05$ . Gyr. = gyrus; SI = primary somatosensory cortex; Postc. = Postcentral; Th. = Thumb; Ringf. = Ring finger.
Group-level contrasts of ring finger and thumb stimulation reveal distinct clusters in the contralateral S1 associated with peak t-values in area 3b and area 1 of S1, respectively. Coordinates given in MNI-space, BA: Brodman area.

**Supplementary Table 3:**
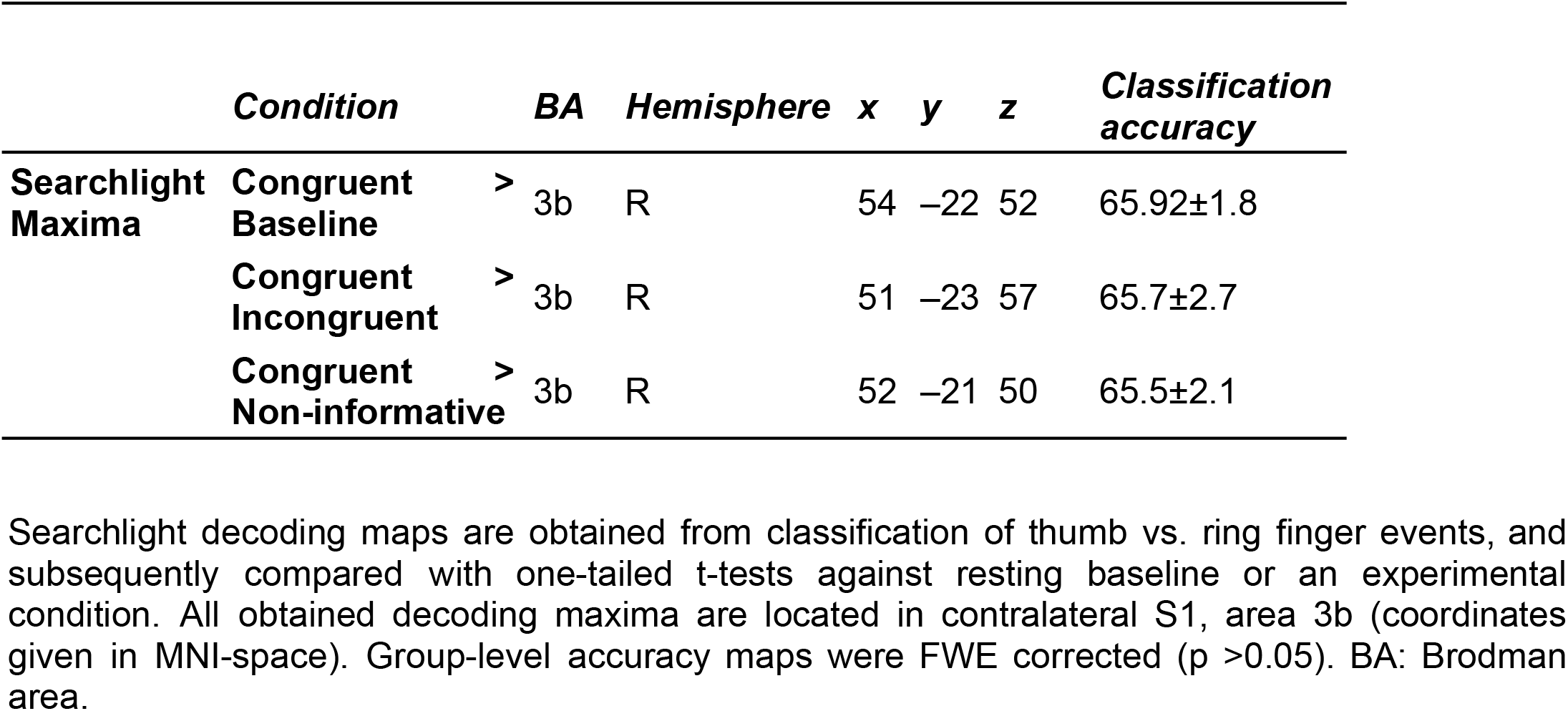
Group-level searchlight maximal accuracies for classification of ring finger versus thumb stimulation.

